# Nest site selection and productivity of a critically endangered parrot, the great green macaw (*Ara ambiguus*), in an anthropogenic landscape

**DOI:** 10.1101/2023.01.10.523381

**Authors:** Thomas C. Lewis, Ignacio Gutiérrez Vargas, Claire Vredenbregt, Mario Jimenez, Ben Hatchwell, Andrew P. Beckerman, Dylan Z. Childs

## Abstract

Nest site selection is the principal way secondary cavity-nesting species mitigate the negative effects of factors such as predation, parasitism and exposure on productivity. High-quality cavities could then be expected to be selected in response to the primary threat to nest success. Understanding how demographic rates are affected by anthropogenic changes to ecosystems is vital if effective conservation management strategies are to be developed and implemented. Large-bodied secondary cavity-nesting birds rely on large cavities in mature trees that are often absent or reduced in anthropogenically disturbed forests. Thus, the availability of high-quality nest sites may be limited for these species, potentially reducing productivity. The aim of this study was to investigate nest-site selection and the effect of nest-site features on productivity in the critically endangered great green macaw (*Ara ambiguus*) in northern Costa Rica. We show that great green macaws select nest sites according to the characteristics of the cavity and of the tree in which they are located. Moreover, productivity was a function of certain cavity features. We conclude that great green macaws are not reliant on primary forest for nest sites and typically choose cavities in remnant, isolated trees in pasture or young secondary forests.

## 1. Introduction

Parrots are one of the most endangered bird families, with 28% of species threatened with extinction globally (IUCN, 2020). They are found throughout the tropics and sub-tropics (Forshaw, 2010; Parr & Juniper, 2010), where the major threats are poaching and habitat loss (Wright et al., 2001; Stojanovic et al., 2016; Berkunsky et al., 2017; Vergara-Tabares et al. 2020), both of which can impact the productivity of a parrot population by reducing the number of good quality nest sites (DeWalt, Maliakal and Denslow, 2003; Amininasab et al., 2016), or by removing nestlings (Wright *et al*., 2001; Hart *et al*., 2016; Martin, 2018). Understanding whether and how the productivity of a parrot population is constrained is essential information when designing a conservation strategy. Without this knowledge, the high financial cost of carrying out conservation (McCarthy *et al*., 2012) may be wasted on ineffective interventions (Schrott, With and King, 2005).

Long-lived species are generally sensitive to reductions in adult survival, so changes in this vital rate are often the main driver of population decline (Sæther & Bakke, 2000). However, this is not always the case in species such as parrots, where nest poaching is a significant driver of the decline in many parrot populations (Wright *et al*., 2001; Hart *et al*., 2016; Martin, 2018). Charismatic and easily accessible species, such as the yellow-naped Amazon (*Amazona auropalliata*), can suffer catastrophic population decline due to poaching of nestlings, with 100% of nests being poached in some areas (IUCN, 2020; Dahlin et al., 2018). As well as poaching, inter- and intra-specific competition can affect productivity by reducing nest site availability (Rendell & Robertson, 1989; Salinas-Melgoza, Salinas-Melgoza and Renton, 2009; Heuck *et al*., 2017), which may then be amplified by habitat degradation reducing the absolute number of available cavities (Cockle et al., 2010; DeWalt et al., 2003). Reduced availability of high-quality nest sites drives individuals to select lower-quality sites, which may cause a decline in population-level productivity (Carrete et al., 2006; Heuck et al., 2017). Thus, it is essential to understand the processes affecting productivity decreases in long-lived species subject to significant impacts (e.g. Tomillo *et al*., 2008; Jepson et al., 2016; Jourdain et al., 2019).

Most parrots are secondary cavity nesters meaning they rely on cavities created by other species to nest in (Forshaw, 2010; Parr & Juniper, 2010). Large-bodied secondary cavity nesters such as parrots are particularly reliant on mature trees that contain cavities of sufficient size (Marsden & Jones, 1997; de la Parra-Martínez *et al*., 2015; Renton *et al*., 2015). The availability of large trees and cavities is limited in degraded and regenerating forests (DeWalt et al., 2003). This suggests that the productivity of secondary cavity-nesting species such as parrots may be limited by nest-site availability in degraded forests (de la Parra-Martínez *et al*., 2015; Renton *et al*., 2015; De Labra-Hernández & Renton, 2016; Stojanovic *et al*., 2021).

Choosing the physical characteristics of nest sites is one of the few ways individuals can counter common threats such as predation (Stojanovic et al., 2017) and parasitism (Tomás et al., 2020). For many parrot species, predation accounts for a significant proportion of nest failure (Berkunsky et al., 2016; Pizo et al., 2008; Renton and Salinas-Melgoza, 2004). Thus, we might expect productivity to be correlated with physical nest-site characteristics that mitigate predation risk (Cockle et al., 2015). Parrots often select cavities that are deeper, higher above the canopy or larger than unused cavities (de la Parra-Martínez et al., 2015; De Labra-Hernández and Renton, 2016; Olah et al., 2014; Saunders et al., 1982; Stojanovic et al., 2021, 2017; Webb et al., 2012). These characteristics have been shown to have a significant positive influence on productivity in the Eclectus parrot (*Eclectus roratus*) (Heinsohn, 2008) and scarlet macaws (*Ara macao*) (Olah *et al*., 2014). This may be because deep cavities offer greater protection against parasites (Tomás *et al*., 2020), or higher cavities have a better field of vision to detect predators (White, Brown and Collazo, 2006). It follows that breeding at low-quality sites is often associated with lower productivity (e.g. Rendell and Robertson, 1989; Stokes & Boersma, 1998; Safran, 2006; Carrate, Donazar and Margalida, 2008). Thus, productivity might be limited in degraded ecosystems if constrained nest-site selection causes parrots to nest in low-quality cavities.

We investigated whether a link between nest-site selection and productivity exists in the critically endangered great green macaw (GGM, *Ara ambiguus*; BirdLife International, 2020). The GGM is a large neotropical parrot whose range extends from the Caribbean lowland forest in Honduras to Colombia and the Pacific coast of Panama down to the dry forest in western Ecuador (Forshaw, 2010; Parr & Juniper, 2010). In Costa Rica, evidence suggests that their steep decline was due to habitat loss; 90% loss of a critical nesting and food tree species, the mountain almond (MA - *Dipteryx panamensis*) (Powell et al., 1999) and a 30% reduction in forest cover to ∼50% by the year 2000 (Calvo-Alvarado et al., 2009). With this evidence, we might expect nest-site availability to limit productivity. Therefore, we aimed to answer two questions:

1. What cavity features do GGMs use to select a nest site?
2. Does the availability of suitable cavities limit productivity?

## 2. Methods

### 2.1 Study site

The study area is a ∼1000km^2^ fragmented Caribbean lowland forest region situated in northern Costa Rica within the San Juan La Selva biological corridor (Fig. 1). Land use is split between cattle pasture, pineapple and other annual crops, and primary and secondary forests (Fagan et al., 2013; Jadin et al., 2016). Annual rainfall is ca. 4667mm (2009-2014), with a slightly drier period between January and April (Gilman et al., 2016) which corresponds to the breeding season of the GGM (Monge *et al*., 2003, 2012).

**Figure 1:**
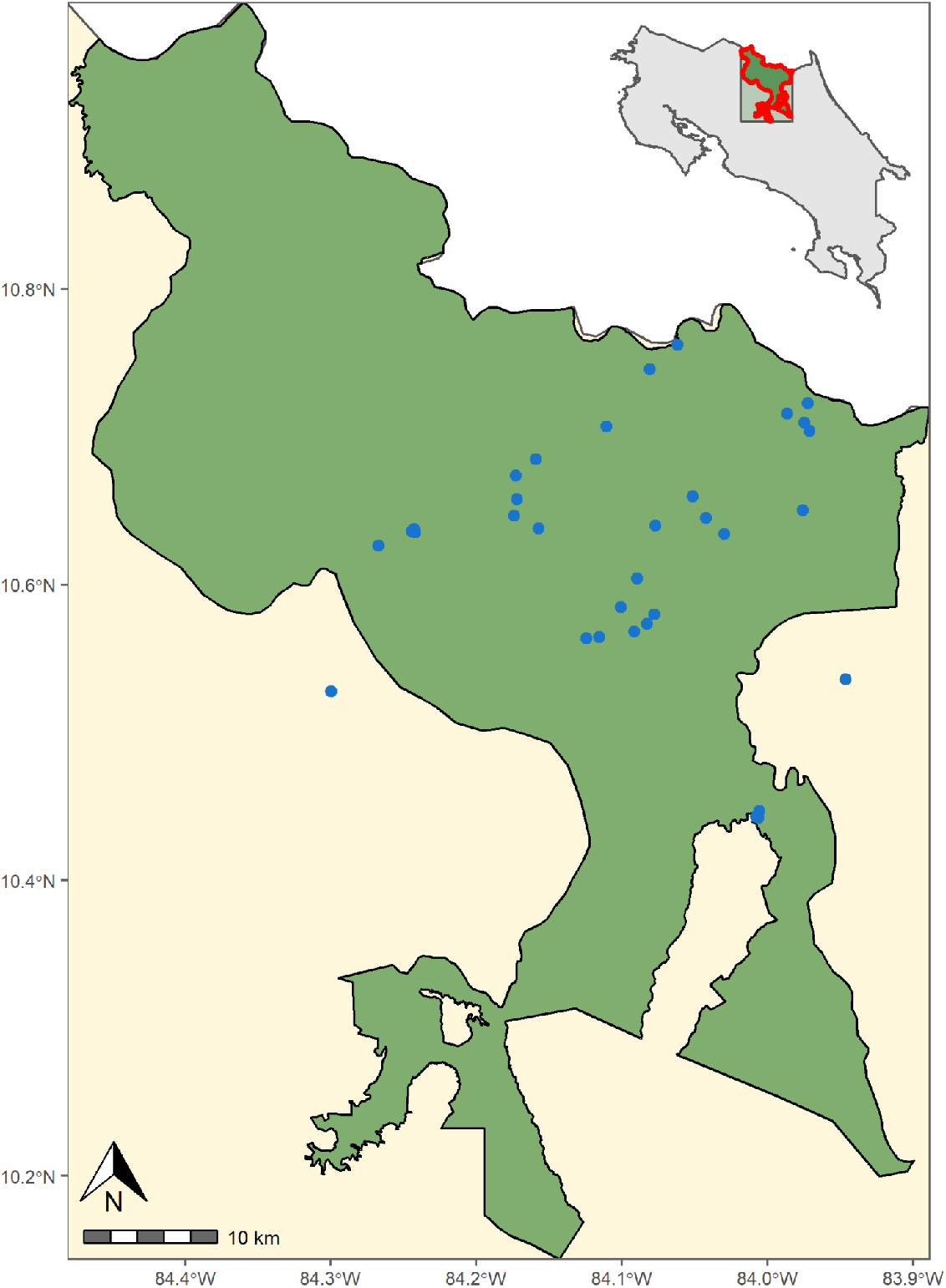
Study site map Study site in the Caribbean lowlands within the San Juan La Selva (SJLS) biological corridor in northern Costa Rica, with nest cavities (blue) marked. There were two monitored nests outside the SJLS biological corridor.

### 2.2 Cavity surveys

The locations of nest cavities come from a database of GGM nest sites collated by Centro Cientifico Tropical (CCT: 1995 - 2020) and Macaw Recovery Network (MRN: 2017 – 2020). Around each active nest cavity (hereafter, a ‘focal cavity’) in the 2020 breeding season between Dec 2019 and May 2020, we surveyed a 500m radius with a Mavic Pro Zoom drone to search for visible cavities. The drone was flown in transects 100m apart and searched for emergent trees. We focused on emergent trees as other studies have found macaws select these types of trees in tropical forests (Renton and Brightsmith, 2009), most likely due to difficulties accessing cavities within a closed canopy forest. We defined an emergent tree as one whose crown was entirely above the surrounding canopy. Once located, it was circled to attempt to detect any cavities; if we found a cavity, a picture was taken directly above the tree. The geo-location on the image was then entered into the GPS so the climbing team could navigate back to the tree on the ground, who then climbed the geo-tagged trees to measure all its cavities.

### 2.3 Features

We took cavity measurements to calculate the entrance area, internal circumference and cavity depth. As cavity entrances are not commonly circular, we treated each entrance as five separate shapes to get the most accurate approximation of the area (Supporting Online Information Fig. S1 & S2). Cavity depth from the lip is the distance from the bottom lip of the cavity entrance to the cavity floor. We calculated internal circumference using cavity width and breadth at the level of the cavity lip. Internal circumference was used instead of measuring the cavity floor because, in many cases, measuring this directly was impossible as the cavities were too deep. We also recorded the cardinal direction of the entrance.

The tree-level features measured were species identity, cavity height from the ground (metres), tree circumference at breast height (metres), canopy connectivity (%) and vertical distance to the canopy to the nearest 5m (metres). We estimated the mean canopy height within a 50m radius of each cavity using the dataset provided by Lang et al. (2022). This global dataset of mean canopy height at 10m resolution was created using deep learning trained with GEDI and Sentinel-2 satellite data. We also estimated tree cover within 50m using the dataset developed by Karra *et al*. (2021); this is a global dataset of landcover/land use created using deep-learning and Sentinel-2 data and trained on 5 billion human-labelled pixels (S3).

### 2.4 Productivity

We monitored productivity during the breeding season of 2020 (Dec 2019 – June 2020). Productivity was measured as the number of chicks fledged per breeding attempt, estimated as the number of fully feathered chicks last seen in the nest before identifying an empty nest. We defined a breeding attempt as a clutch of eggs reaching a conclusion, either success or failure.

### 2.5 Imputation

We could not acquire complete measurements for every cavity due to cavities being challenging to access or human error where measurements were missed in the field. We imputed these missing data to maximise the power of our statistical analyses. We used multivariate imputation by chained equations (mice); this is a more robust way to deal with incomplete data sets compared to single imputation or data deletion approaches (Penone et al., 2014; Taugourdeau et al., 2014; Cooke et al., 2019). We used the mice package to generate our imputed values (Buuren and Groothuis-Oudshoorn, 2011). We used a random forest imputation method, with all numeric variables to impute missing values. Finally, we imputed 100 datasets to capture the uncertainty in the imputation process.

### 2.6 Statistical analysis

#### Cavity feature selection

To determine whether GGMs select cavity features, we fit a pair of intercept-only Bayesian regression models using the default flat prior. For each feature *i* to model the distribution of its values in occupied 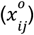 and unoccupied 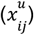 cavities:

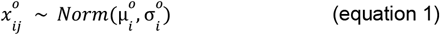

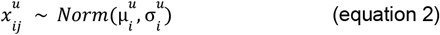

Where 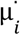 and 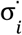 are the mean and standard deviation of their distributions. We compare features of occupied and unoccupied cavities by calculating the posterior distribution of the difference in their means and standard deviations:

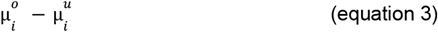

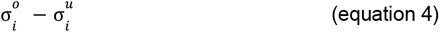

If the 95% credibility intervals of the differences did not cross zero, we treated the feature as selected for. By evaluating both standard deviation and mean, we can approach selection in different ways. The difference in the means and their distributions tells us whether they are actively selecting these features. We expect the direction of the difference between means will vary depending on the cavity feature, for example, the difference would be positive if larger entrance holes are preferred compared to what is available. Whereas standard deviation tells us how specific they are with their selection of a feature. For this, we would expect strongly selected features to have a negative difference in standard deviation because occupied cavities would have a smaller standard deviation than unoccupied ones.

#### Nest suitability

To evaluate the suitability of each occupied cavity, we then calculate the cavity suitability scores for each feature (*s*_*ij*_) that differs among the occupied and unoccupied cavities:

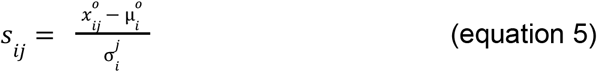

This gives the standardised score 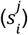 for feature *i* associated with cavity *j*; the further away from the mean or suitable value the lower the score will be. Therefore less suitable cavities have lower scores. We combined these standardised scores into the nest suitability score (NSS) by taking the mean of the selected features.

Using the NSS, we carried out an ordinal regression with a logit link and a flexible threshold to explore the association with this integrative measure. We then compared the performance of each model using the Widely Applicable Information Criterion (WAIC - Watanabe, 2010). We then used the top-performing model to estimate how many cavities not used by GGMs are within the nest suitability score range and, therefore, suitable for GGM use.

We used the `brms` package to run all of our Bayesian models; this permitted the pooling of results across imputed datasets and examination of posterior distributions (Bürkner, 2021, 2018, 2017), which is crucial to our analysis. We carried out all analyses in R (R Core Team, 2021).

## 3. Results

We measured a total of 192 cavities; 95 were found using a drone, 66 were from an existing Centro Cientifico Tropical (CCT) database, and 51 had been found incidentally by the field team of Macaw Recovery Network. We found an additional 83 trees in the drone survey but did not include them in the study due to aggressive bees (n = 2), no cavities being found in the marked tree (n = 2), and fieldwork being cut short by the global Covid-19 pandemic (n = 79). We excluded 22 cavities that were in entirely hollow trees. There was an average of 2.65 cavities per tree (n = 192). We found six species of trees with cavities; five were used as nest trees (Fig. S6). Most located cavities were in mountain almonds (88.5%; n = 185), as were the majority of nests (84.6%; n = 33). These results indicate that GGMs use mountain almonds because this species is the most abundant source of cavities rather than actively preferring the species.

### 3.1 Cavity feature selection

The difference in the posterior distributions of the mean cavity features in occupied and unoccupied cavities suggest that GGMs select for entrance height (mean = -0.69, 95% CI = -1.06 to -0.33), cavity depth (mean = 0.39, 95% CI = 0.034 to 0.749), internal circumference (mean = 1.14, 95% CI = 0.83 to 1.46), and entrance area (mean = 0.56, 95% CI = 0.23 to 0.90). At the tree level, they use canopy connectivity (mean = -0.54, 95% CI = -0.93 to -0.19) when selecting nest cavities, as well as canopy height (mean -0.45, 95% CI = 0.-83 to -0.07) and tree cover (mean = -0.46, 95% CI = -0.87 to -0.04) at the local area level (Fig. 2A & C). Whereas the difference in posterior distributions of the standard deviation shows that GGMs have a narrower preference for cavity entrance area (mean = -0.37, 95% CI = -0.61 to -0.19), depth (mean -0.28, 95% CI = 0.53 to -0.01) and canopy connectivity (mean = -0.79, 95% CI = -1.13 to -0.49) (Fig. 2B & D).

**Figure 2:**
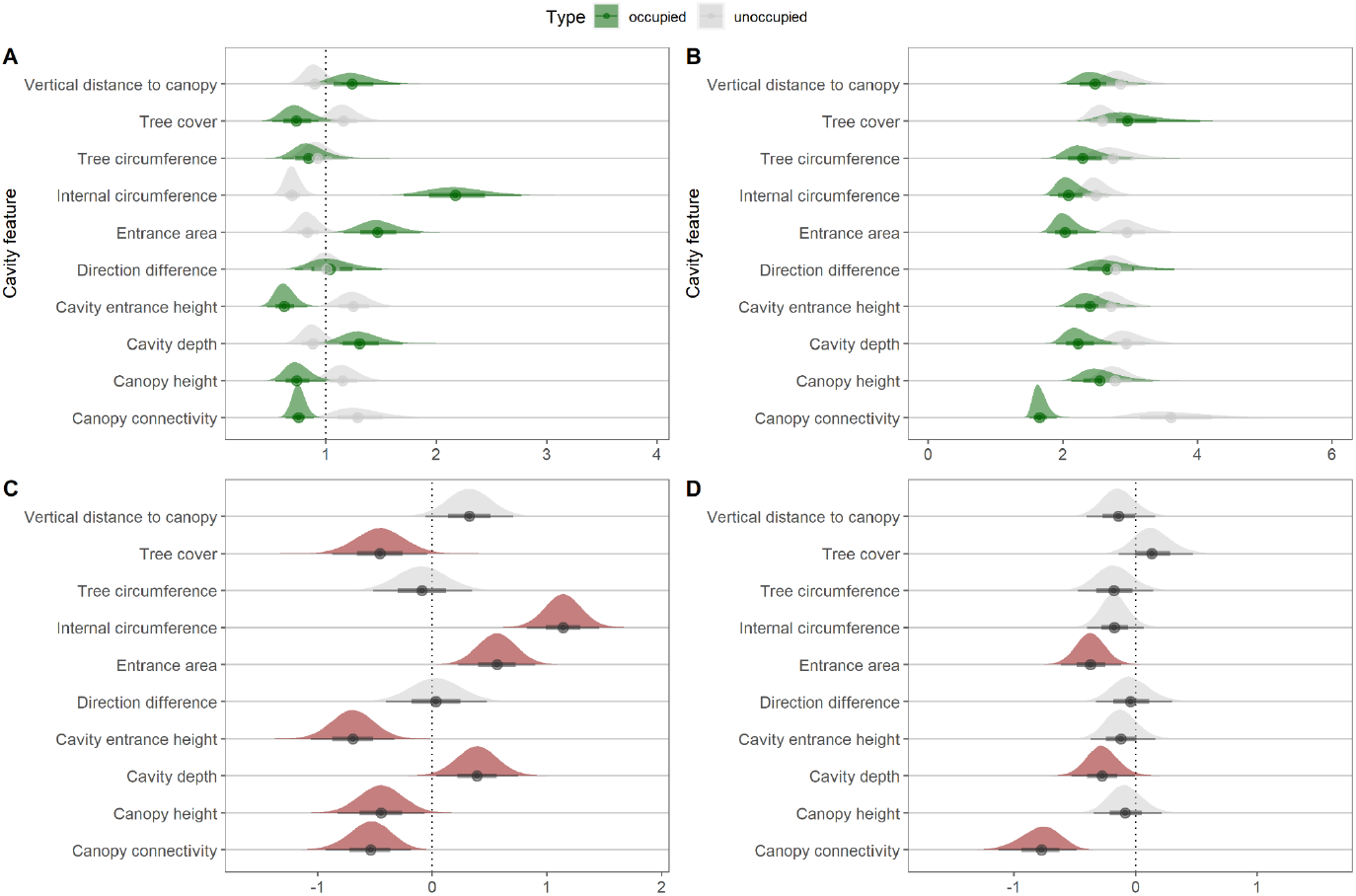
Univariate model outputs for cavity characteristics Univariate model A) means and posterior distributions and B) variance and posterior distributions for occupied and unoccupied cavities. With the difference in C) means and posterior distributions and D) standard deviation and posterior distributions for each cavity feature with significant features, those with 95% credibility intervals not crossing zero, are highlighted in red.

### 3.2 Nest suitability

During the 2020 breeding season, 47 GGMs fledged from 35 breeding attempts, a mean of 1.34 fledglings per breeding attempt. The maximum number of chicks fledged was 3 (n = 1). When run individually, entrance area (0.23, 95% CI = -0.94 to 1.44) and internal circumference (0.62, 95% CI = -0.11 to 1.42) were positively correlated with the number of fledglings per breeding attempt, but only cavity depth (1.05 95% CI = 0.20 to 1.99) was significant (Fig 3). Cavity entrance height (−0.17, 95% CI = -0.90 to 0.53) and canopy connectivity (−0.59, 95% CI -2.22 to 0.96) were negatively correlated with the number of fledglings, but the relationships were not significant. Nest suitability score (2.20, 95% CI = 0.33 to 4.34) was significantly positively correlated to the number of fledglings per breeding attempt.

**Figure 3:**
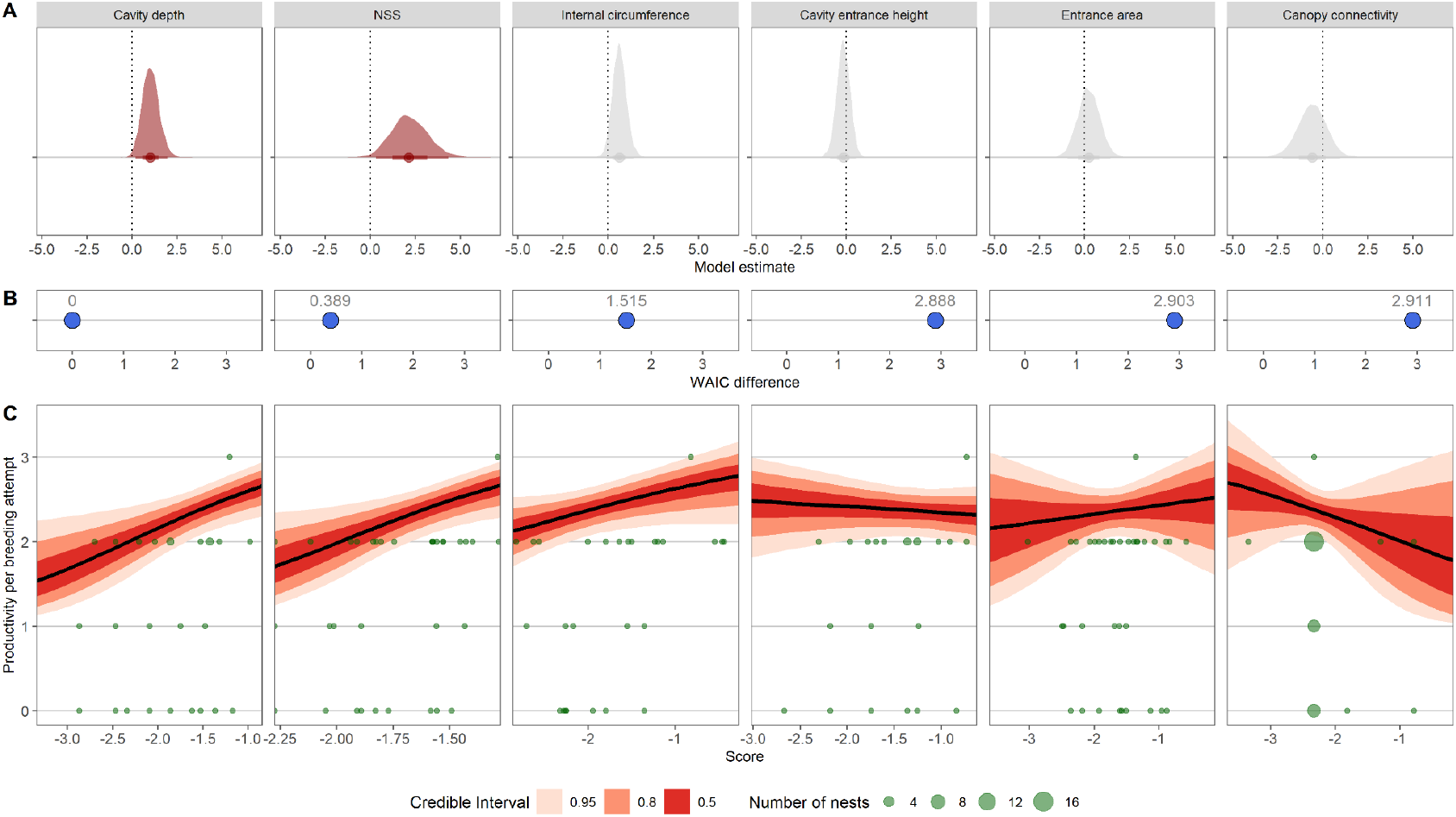
The relationship between productivity and cavity features. A) Model estimates of the regression slopes and their posterior distributions from the univariate ordinal regression of each cavity feature, with the significant models in red. B) The difference in WAIC scores compared to the best-performing model. C) The relationship between each feature and predicted productivity alongside the data (green).

## 4. Discussion

The GGMs in our study population prefer larger, deeper cavities in isolated trees, showing a strong selection for larger cavity entrance area, greater depth and low canopy connectivity. This agrees with numerous other studies that found that parrots select for cavity depth (de la Parra-Martínez et al., 2015; De Labra-Hernández and Renton, 2016; Olah et al., 2014; Renton et al., 2015; Saunders et al., 1982; Stojanovic et al., 2021, 2017; Webb et al., 2012) or tree isolation (Berkunsky et al., 2016; Britt et al., 2014; Renton et al., 2015). We know that predation accounts for a significant proportion of nest failure in many parrot species (Pizo et al., 2008; Renton et al., 2015; Berkunsky et al., 2016); therefore, the selection of these specific features is likely driven by the need to reduce predation risk. By selecting isolated trees, parrots can passively reduce predation risk from nonvolant predators (Berkunsky et al., 2016; Britt et al., 2014; de la Parra-Martínez et al., 2015), whilst deeper cavities reduce the risk from avian predators (de la Parra-Martínez et al., 2015; Mejías et al., 2017; Wesolowski, 2002; Zhu et al., 2012).

For secondary cavity-nesting species, cavity location and morphology are the only way to reduce predation risk passively. Therefore, the relationship between cavity and tree characteristics and productivity can indicate which type of predators are the current primary threat to GGM nesting success. Although it simplifies the suite of factors such as habitat quality (Dhondt, 2010; Jones et al., 2014) and climatic conditions (Borgman and Wolf, 2016; McGillivray, 1981) that influence productivity, it is a valuable process as it might highlight potentially information for conservation managers. Birds can attempt to reduce the risk from avian predators by selecting deeper cavities (Wesolowski, 2002; Zhu et al., 2012).

Therefore, if avian predation was the primary cause of nest failure, we might expect to see a relationship between cavity depth, as birds are more likely to predate shallow nests (Mejías et al., 2017). Alternatively, individuals passively reduce accessibility to nonvolant predators such as snakes and arboreal mammals by selecting isolated trees because they need access to cavities via connected canopy or vines (Berkunsky et al., 2016; Britt et al., 2014; Koenig, 2001). Therefore, if nonvolant predators are the primary source of nest predation, we expect to see a relationship between canopy connectivity and productivity. We have found a significant positive relationship between productivity and cavity depth, suggesting avian predation is one of the primary drivers of nest failure. Indeed, we have direct evidence of this in our study population, where five of seven confirmed predation events from our study area were due to avian predators (MRN *unpub. data*). Therefore, for conservation managers focusing on addressing the threat from avian predators may be a cost-effective way to increase nest success and boost productivity.

GGMs are apparently unique in their selection of cavities with entrance heights lower than surrounding cavities (de la Parra-Martínez et al., 2015; Renton and Brightsmith, 2009). This may be an adaptation to fragmentation and degradation of habitat in this area of Costa Rica (Chassot et al., 2007; Grantham et al., 2020; Karra et al., 2021), where large cavities in large, emergent trees in the forest have likely been lost (Chassot et al., 2007; DeWalt et al., 2003). Cavity size is inversely related to height in the tree (Lindenmayer et al., 2000); therefore, low tree cover and canopy height could make large cavities lower down in trees accessible to GGMs. This might partially compensate for the loss of mature emergent trees in this region over the last 100 years (Sader and Joyce, 1988). We monitored two active nests in 2020 that were 9.4m and 10.4m above the ground. The average canopy height across the San Juan La Selva biological corridor is 23.4m (Lang et al., 2022), suggesting that if these trees were within the forest, they would not be accessible or suitable for GGMs.

Interspecific competition can affect productivity by limiting the number of available cavities (Berris et al., 2022; Bonaparte and Cockle, 2017; Symes and Perrin, 2003). Half of the cavities occupied by interspecific competitors were used by scarlet macaws (Supporting Online Information Table S1). Research has shown that scarlet macaws have a broad nesting niche and compete directly with other large macaws for nesting cavities (Renton and Brightsmith, 2009). In our study, the cavities occupied by SCMs are less suitable for GGMs, suggesting that there is either limited competition for cavities or GGMs are stronger competitors. However, as suitable cavities remain unoccupied and both the SCM and GGM populations in this area are recovering, competition may increase in the future.

### 4.1 Conservation implications

Cavity entrance height has often been found to be an important factor in nest site selection for macaws, with species selecting for cavities higher than unused cavities (de la Parra-Martínez et al., 2015; Olah et al., 2014). However, we found that GGMs nest in cavities lower than unused cavities. This may be due to the lack of forest around nest trees. Costa Rica reached a 17% forest cover low in the 1980s (Sader and Joyce, 1988), and although logging continues in this region, large-scale deforestation no longer occurs (Fagan et al., 2013). In many areas, regenerating forest has replaced cattle pasture (Fagan et al., 2013; Jadin et al., 2016). However, as the forest continues to recover, there is a risk that cavities currently occupied by GGMs will become more connected to adjacent trees, increasing the accessibility for nonvolant predators. This is a natural process that we have seen at a few nests in abandoned pastures; when the nests were first monitored 20 years ago, they were in isolated trees, and now they are within the canopy (Aleman *pers comms*).

Trees around nest sites could be managed to maintain limited connectivity with other trees and prevent access to the canopy for nonvolant predators. However, installing artificial nest boxes may represent a better management option than preventing forest recovery around current nest trees. Nestbox provision is a common conservation strategy to combat limited cavity availability in degraded habitats (Berthier et al., 2012; Darling et al., 2004; Jones et al., 1995; Tollington et al., 2013). For example, nest boxes have been successfully employed with another critically endangered macaw, the blue-throated macaw (*Ara glaucogularis*) in Bolivia (Herzog *et al*., 2021). Placement and design for GGMs could utilise the isolated trees whilst allowing for a design that reduces the risk of avian predation. A secondary benefit is that it would permit easier monitoring and active management of breeding attempts if necessary (Jones et al., 1995; Tollington et al., 2013). A small experiment with four nestboxes of one design was carried out by the Centro Cientifico Tropical (CCT) between 2016 and 2020; one chick successfully fledged from a nestbox in 2020 (CCT *unpub. data*). This demonstrates that wild GGMs will use nest boxes and suggests that further supplementation could be a viable management strategy. However, using next boxes as a management tool does come with associated costs. It would have to be a long-term strategy to be able to mitigate the loss of cavities while others develop in maturing trees.

## 5. Conclusion

Habitat loss and degradation have caused the loss of mature trees across the tropics, resulting in the loss of large cavities for large-bodied cavity nesters such as the GGM (DeWalt et al., 2003; Degen et al., 2006; Cockle et al., 2010). As anthropogenic disturbance in tropical ecosystems grows, we need to understand how endangered species are affected and adapt to their new environment. Ecological traps can form if cues are used to select suitable nest sites, and actual nest site quality become uncoupled due to anthropogenic disturbance. This can lead to declines in productivity and subsequent population decline (e.g. Tozer et al. 2012; Zhu et al. 2012; Díaz and Kitzberger 2013). It is encouraging that we found a connection between nest-site selection and productivity in this anthropogenically altered landscape (Chassot et al., 2007; Grantham et al., 2020; Karra et al., 2021), suggesting that no ecological trap has formed. By studying nest-site selection and productivity together, we have identified factors that could potentially limit the future recovery of the critically endangered GGM. Finally, we suggest that nestbox provisioning could be a solution to the potential loss of nest sites to forest regeneration. This approach can be successful but does mean a long-term commitment to maintenance for conservationists.

## Acknowledgements

This study was funded through the National Environmental Research Council’s (NERC) ACCE Doctoral Training program (NE/L002450/1) with support from Macaw Recovery Network, Costa Rica and Chester Zoo, UK. All work as carried out in Costa Rica was done so under the permit R-SINAC-PNI-ACAHN-29-2019

Nest site selection and productivity of a critically endangered parrot, the great green macaw (Ara ambiguus) in an anthropogenic landscape

## 7. Figures and tables

**Table 1:**
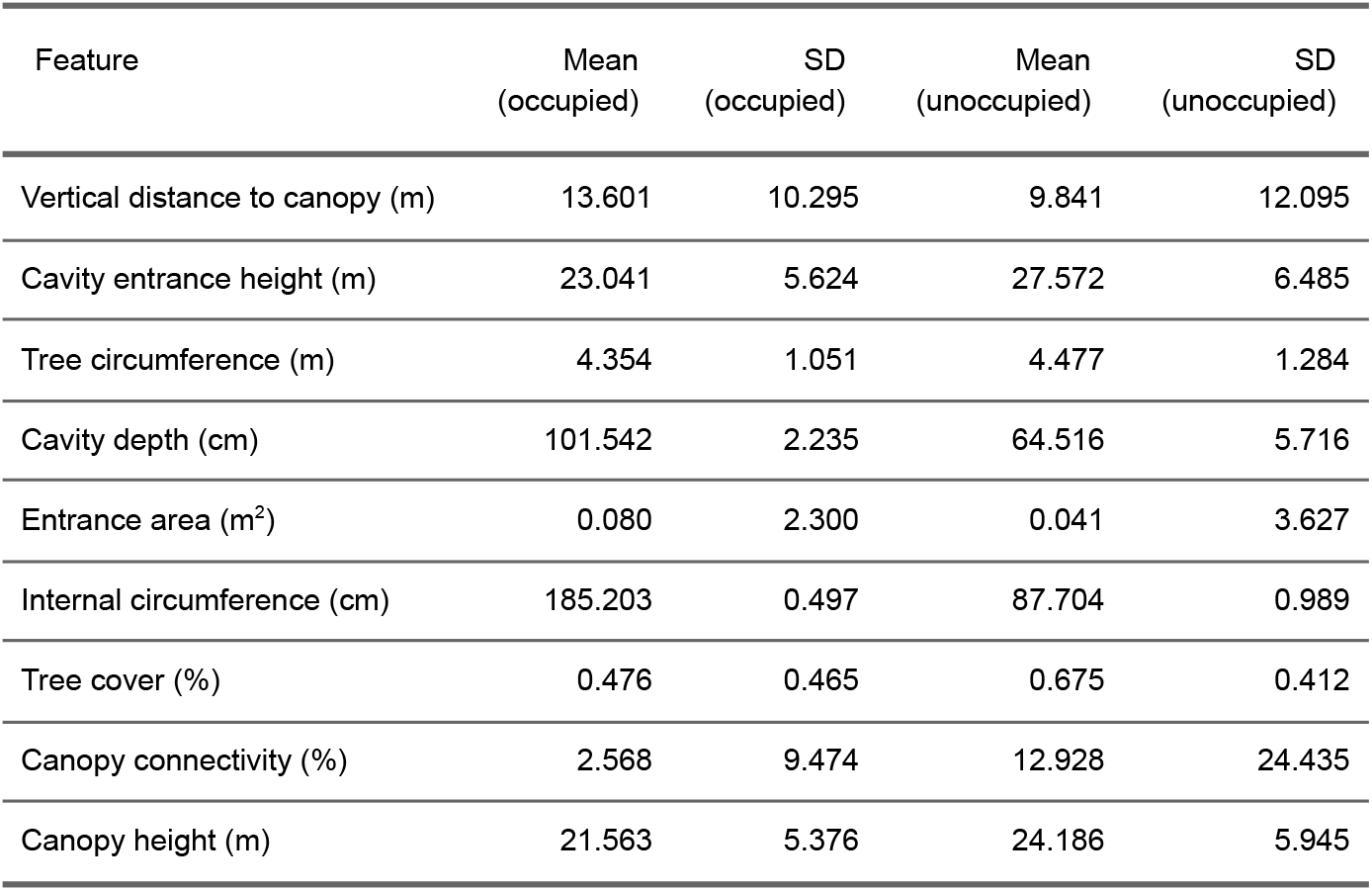
Cavity characteristics Summary of all tree and cavity characteristics of occupied (n = 37) and unoccupied (n = 79) cavities.

## Notes

### Competing Interest Statement

The authors have declared no competing interest.

