## Supplementary material for "Nest site selection and productivity of a critically endangered parrot, the great green macaw (*Ara ambiguus*), in an anthropogenic landscape": Online Supplementary Materials

### Figure S1:

Cavity entrance diagram. a) Three cavity entrance measurements were taken in the field, which allowed b) an approximate calculation of the cavity entrance area by splitting the cavity into four shapes.

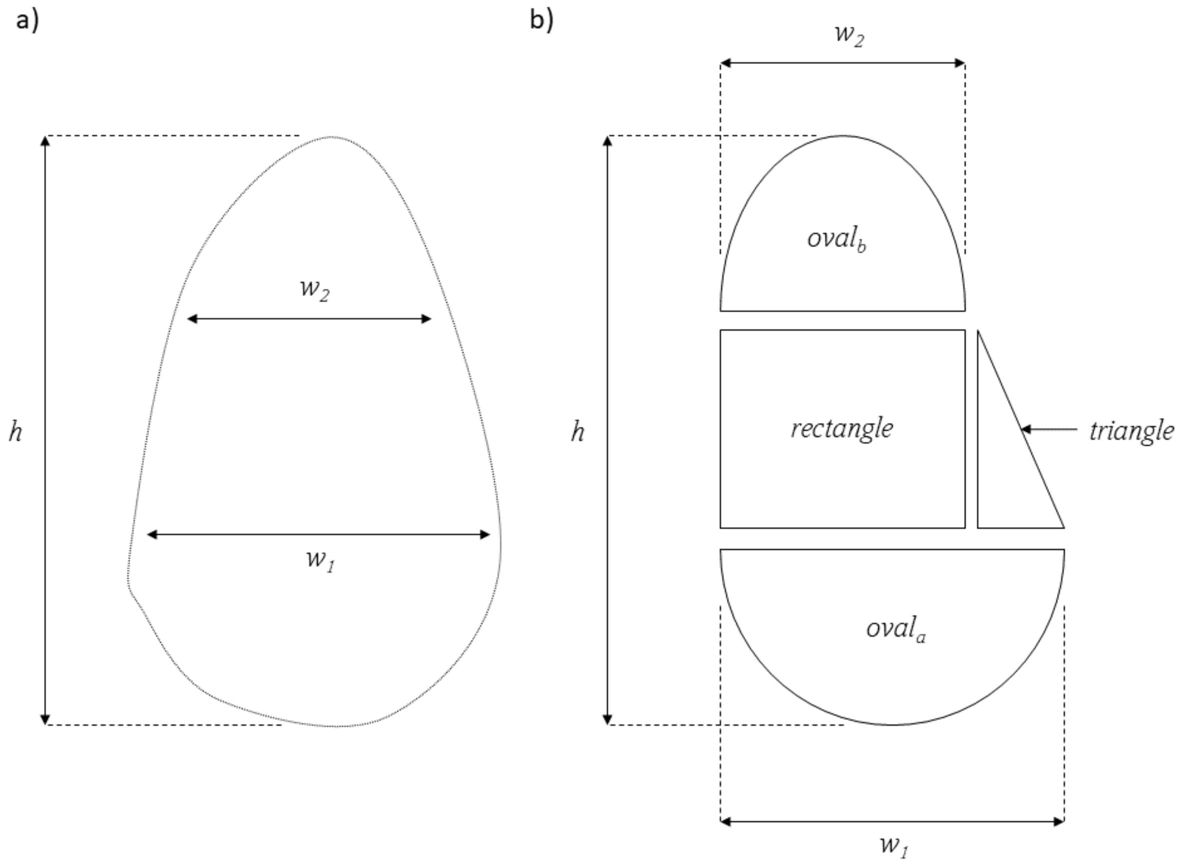

### Equations S1:

Entrance area equation:

$$oval_a = \frac{\frac{h}{3} \cdot \frac{w_1}{2} \cdot \pi}{2} \quad \text{(equation 1)}$$

Where $h$ = cavity entrance height (cm) and $w_1$ = cavity width at ⅓ of cavity entrance height (cm).

$$oval_b = \frac{\frac{h}{3} \cdot \frac{w_2}{2} \cdot \pi}{2} \quad \text{(equation 2)}$$

Where $w_1$ = cavity width at ⅔ of cavity entrance height (cm).

$$rectangle = min(w_1, w_2) \cdot \frac{h}{3} \quad \text{(equation 3)}$$

$$triangle = \frac{max(w_1, w_2) - min(w_1, w_2) \cdot \frac{h}{3}}{2} \quad \text{(equation 4)}$$

$$area = \frac{rectangle + triangle + oval_a + oval_b}{10000} \quad \text{(equation 5)}$$

Where $area$ is in $m^2$

### Equations S2:

Internal circumference equation

$$i = \pi \cdot \sqrt{2 \cdot (0.5 \cdot w)^2 + (0.5 \cdot b)^2} \qquad \text{(equation 6)}$$

Where $i$ is the internal circumference.

Figure S2:

Visualisation of a correlation matrix between all cavity features. We transformed three skewed features: cavity depth, entrance area and internal circumference. There are a number of highly correlated features, but as we are running univariate models of each we did not remove any from our analysis (* = 0.05, ** = 0.01, *** = 0.001).

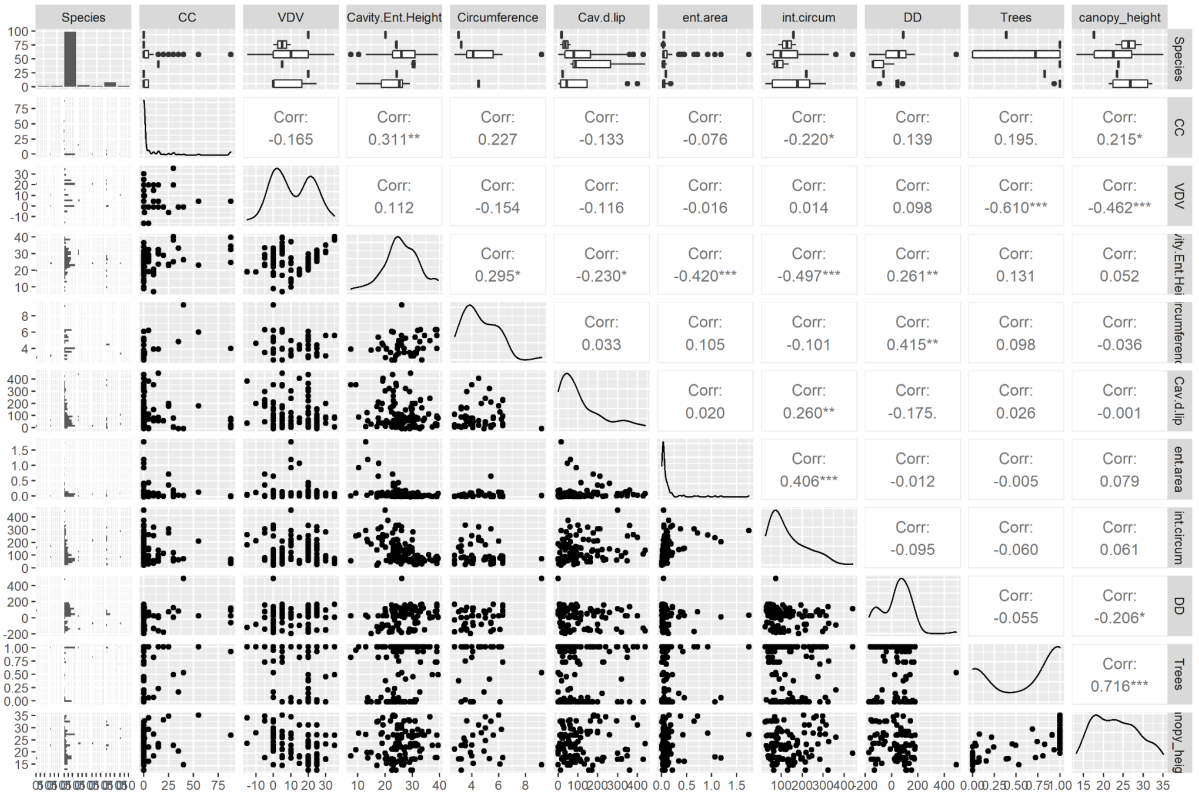

Figure S3:

Species of trees that were occupied and unoccupied showing there is no strong selection for any particular species.

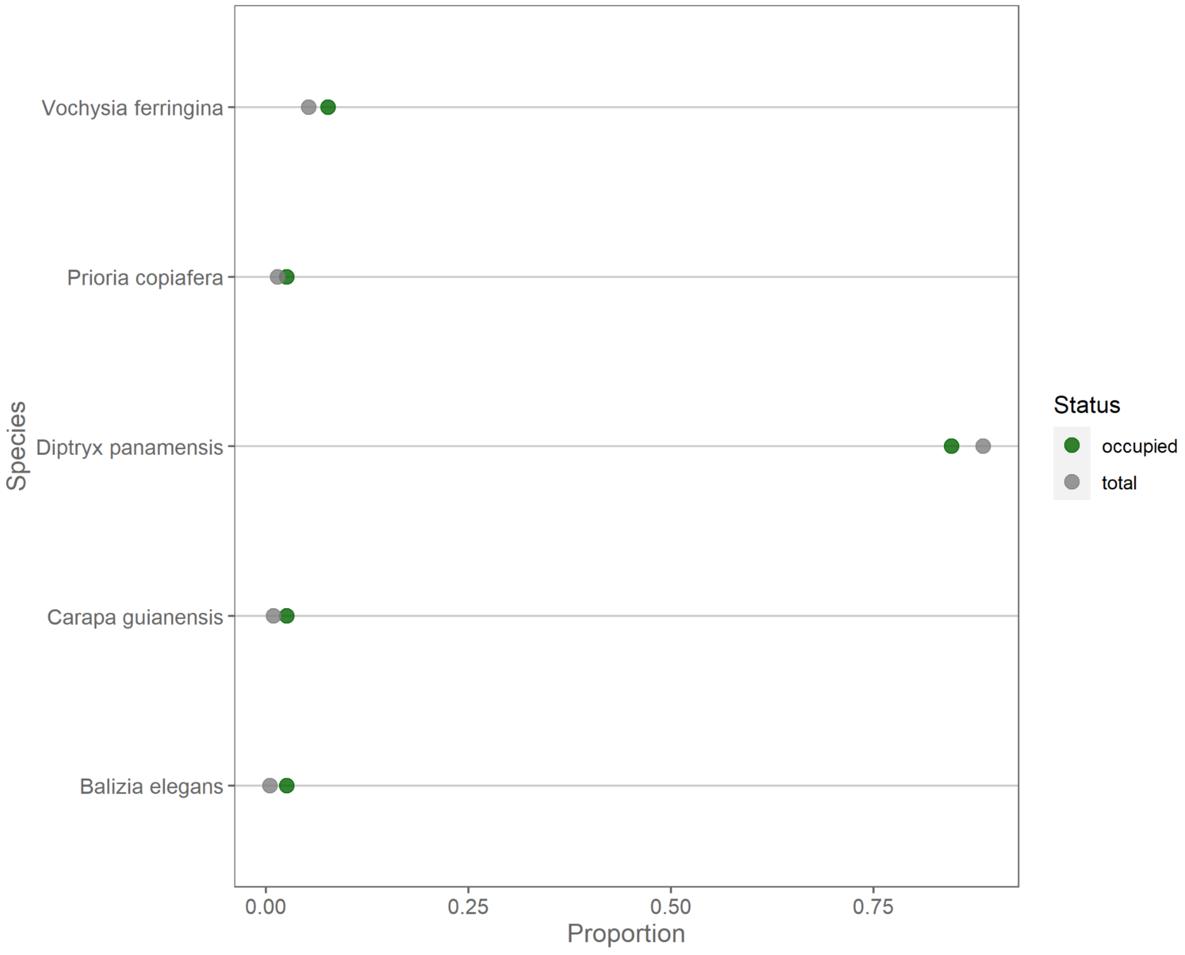

Table S1:

Nest suitability and cavity depth model predictions for all cavities. The two most numerous interspecific competitors *Ara macao* and *Amazona autumalis* have similar predicted productivity by NSS but not cavity depth.

| Species | Number of cavities | Mean productivity | |
|---|---|---|---|
| | | Cavity depth | NSS |
| *Amazona autumnalis* | 5 | 1.39 | 0.97 |
| *Ara ambiguus* | 37 | 1.40 | 1.26 |
| *Ara macao* | 9 | 1.10 | 0.97 |
| Interspecific competitors | 5 | 1.55 | 1.43 |
| unoccupied | 91 | 1.05 | 0.89 |

### Table S2:

Summary of all tree and cavity characteristics of occupied (n = 37) and unoccupied (n = 79) cavities.

| Feature | Mean (occupied) | SD (occupied) | Mean (unoccupied) | SD (unoccupied) |
|---|---|---|---|---|
| Vertical distance to canopy (m) | 13.601 | 10.295 | 9.841 | 12.095 |
| Cavity entrance height (m) | 23.041 | 5.624 | 27.572 | 6.485 |
| Tree circumference (m) | 4.354 | 1.051 | 4.477 | 1.284 |
| Cavity depth (cm) | 101.542 | 2.235 | 64.516 | 5.716 |
| Entrance area ($m^2$) | 0.080 | 2.300 | 0.041 | 3.627 |
| Internal circumference (cm) | 185.203 | 0.497 | 87.704 | 0.989 |
| Tree cover (%) | 0.476 | 0.465 | 0.675 | 0.412 |
| Canopy connectivity (%) | 2.568 | 9.474 | 12.928 | 24.435 |
| Canopy height (m) | 21.563 | 5.376 | 24.186 | 5.945 |
